# Design, Monitoring, and Application of a Cost-Effective Bioreactor for Biogas Production

**DOI:** 10.1101/2025.02.03.636301

**Authors:** Angela Ventura-Prados, Javier Solís-García

## Abstract

This study presents the design and construction of a low-cost bioreactor for biogas production, aimed at regions with abundant organic waste to promote renewable energy use in rural and agricultural communities. The bioreactor is equipped with sensors to monitor critical parameters such as temperature, pH, and Total Dissolved Solids (TDS) to ensure efficient anaerobic digestion, using horse manure as the substrate. Operational conditions were evaluated, maintaining pH between 6 and 7, the optimal range for biogas production, and temperature between 30 and 35°C, ideal for methanogenic activity. TDS values fluctuated between 620 and 780 mg/L, revealing a direct relationship with biogas production and emphasizing the importance of maintaining optimal solid concentrations, as excessive solids reduced efficiency. This work demonstrates the feasibility of implementing low-cost bioreactors for biogas generation in rural areas, contributing to sustainable energy recovery and effective organic waste management.

## 1. Introduction

Anaerobic Digestion (AD) is an efficient and widely recognized biological process for biogas production, involving the degradation of organic matter in the absence of oxygen. This process, carried out by specific microbial consortia, yields biogas as a primary product and a nutrient-rich digestate, which can be used as an agricultural amendment (Hua et al., 2020; Koszel & Lorencowicz, 2015). AD encompasses a series of biochemical, chemical, and physicochemical reactions that require precise control of key operational parameters, such as temperature, pH, mixing, Carbon-to-Nitrogen (C:N) ratio, and Hydraulic Retention Time (HRT) (Wang et al., 2018; Amani et al., 2010).

The AD process occurs in four sequential stages: hydrolysis, acidogenesis, acetogenesis, and methanogenesis. Each stage depends on specialized microorganisms and specific operational conditions (Amani et al., 2010). Methanogenesis, the final and critical phase for biogas production, is particularly sensitive to environmental factors such as oxygen absence and specific temperature ranges—either mesophilic (approximately 37°C) or thermophilic (approximately 55°C). These conditions significantly influence both reaction rates and process stability (Kalaiselvan et al., 2022).

The quality and quantity of biogas produced are influenced by the feedstock characteristics and the operational conditions of the biodigester (Abubakar, 2022). Typically, biogas comprises 60–70 % methane, 20–30 % carbon dioxide, and trace amounts of other gases. Consequently, biodigester design plays a critical role in controlling and optimizing the AD process, ensuring an optimal environment for microbial activity (Kalaiselvan et al., 2022).

This study proposes the construction of a Continuously Stirred Tank Reactor (CSTR) with semi-continuous feeding, designed to maximize process efficiency, reduce costs, and enable implementation in rural areas, thereby promoting the energy utilization of organic waste (Ma et al., 2015). In particular, this work addresses the following objectives:

- Design and construction of an economical and accessible CSTR with semi-continuous feeding, incorporating low-cost materials and technologies to optimize anaerobic digestion.
- Implementation of a real-time monitoring system for key parameters such as temperature, pH, and TDS to ensure stable and efficient operation.
- Evaluation of the influence of operational parameters on biogas production and analysis of the reduction of key contaminants (e.g., nitrogen, phosphorus) in the effluent.

By addressing these objectives, this study demonstrates the feasibility of implementing low-cost bioreactors for sustainable energy recovery and waste management in rural areas. The proposed system combines simplicity, efficiency, and accessibility, making it highly scalable for agricultural and livestock applications.

## 2. Background

Biodigesters are systems specifically designed to optimize the anaerobic digestion process, enabling the conversion of organic matter into biogas through design strategies that maximize efficiency (Kalaiselvan et al., 2022). Based on feeding regimes, these systems can be classified into batch, semi-continuous, and continuous systems, each with specific applications:

### Batch Systems

These systems involve the complete initial loading of substrate into the biodigester, followed by a digestion period without further addition or removal of material. Commonly used in research and small-scale operations, this approach is ideal for assessing the biodegradability of specific residues. However, batch systems require downtime between cycles for emptying and cleaning the biodigester, limiting continuous biogas production (Abubakar, 2022; Xing et al., 2020).

### Semi-Continuous Systems

In this regime, the biodigester periodically receives substrate additions while an equivalent fraction of digestate is removed, maintaining a constant volume. This approach is suitable for small-to medium-scale operations that require a regular, though not uninterrupted, flow of biogas. Continuously stirred tank reactors are the most commonly employed design for this regime due to their versatility and effectiveness in digesting diverse organic materials. Widely utilized in agricultural sectors, CSTRs provide efficient waste management while being adaptable to local energy demands (Lansing et al., 2010; Ma et al., 2015).

### Continuous Systems

These systems are designed for uninterrupted operation, allowing constant substrate addition and simultaneous digestate removal. They are ideal for large-scale industrial and agricultural plants where a steady and maximized biogas production is essential. Upflow Anaerobic Sludge Blanket reactors are the most prevalent in this category due to their high level of automation and precise control over parameters such as pH, temperature, and mixing (Tagne et al., 2021).

Given the high costs of conventional biological reactors and the growing interest in implementing them in agricultural areas for organic waste utilization (Romero-Güiza et al., 2021), this study presents the construction of a CSTR with a semi-continuous feeding system. The design includes an inlet and outlet system for substrate control, a stirring mechanism to ensure homogeneous mixing of organic matter, and a biogas collection system using purpose-designed storage bags (Ahmed et al., 2016). Additionally, low-cost technology has been integrated for monitoring key parameters, such as temperature, TDS, and pH, which are critical for evaluating process efficiency (Abubakar, 2022).

The methodology developed in this study offers an economical and accessible solution to promote biogas production in rural areas through the reuse of organic waste (Appels et al., 2008). This comprehensive approach not only facilitates small-scale experimental testing on various feedstocks but also fosters sustainability and energy recovery in agricultural and livestock communities (Lansing et al., 2008). The following sections detail the design, construction, and performance evaluation of this biodigester, highlighting its potential as a key tool for advancing the transition toward a circular bioeconomy.

## 3. Methods and Materials

### 3.1. Biodigester Design

In this study, a biodigester with a maximum capacity of 20 liters was designed, configured as a CSTR and operated with a semi-continuous feeding system. The bioreactor design includes: (1) an inlet system for feedstock and an outlet system for processed digestate, (2) a stirring rod to ensure the homogeneity of the contents, (3) a monitoring screen for the continuous recording of sensor data, and (4) a Tedlar bag for the collection and storage of the generated biogas (Figure 1).

**Figure 1.**
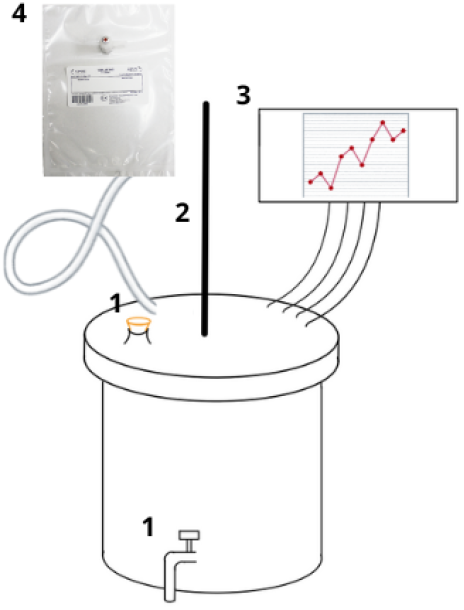
Schematic diagram of the components of the biodigester: (1) inlet and outlet for feedstock and digestate, respectively, (2) stirring rod, (3) monitoring screen for sensor data collection and visualization, and (4) Tedlar bag for biogas collection.

The materials used for the construction and assembly of the biodigester are listed below in Table 1.

**Table 1.** Summary of the quantity and cost of materials used in the construction of the biodigester.

| Materials | Quantity | Unit Cost (€) |
| --- | --- | --- |
| Beer fermentation bucket | 1 | 22.89 |
| Metal rod | 1 | 5.36 |
| Water pipe faucet | 1 | 10.49 |
| Cork stopper | 1 | 6.60 |
| Aquarium heater | 1 | 12.96 |
| Thermal sensor | 1 | 15.30 |
| pH sensor | 1 | 30 |
| TDS sensor | 1 | 26.75 |
| Plastic tube | 1 | 8.79 |
| Rotating motor | 1 | 14.55 |
| Tedlar bag | 1 | 12.68 |
| Silicone | 4 | 4.58 |
| Total cost |  | 156.63 |

### 3.2. Inoculum and Substrate

The inoculum used was prepared together with the substrate, consisting of horse manure obtained from a private agricultural area in Seville. This inoculum was derived from the liquid extracted from a digestate previously produced in the laboratory under anaerobic conditions within a mesophilic range. Prior to its introduction into the biodigester, the horse manure underwent a pretreatment that included thermal treatment and a physical crushing process, both designed to enhance the biodegradability of the material.

The biodigester, with a total volume of 20 liters, was loaded with 16 liters, maintaining 4 liters of free working volume. The mixture of horse manure and inoculum was prepared at a 1:2 ratio (kg of manure/L of inoculum), following the recommendations of previous studies by Romero-Güiza et al. (2021) and Wang et al. (2018). This ratio was selected to optimize the microbiological interaction between the solid substrate and the liquid inoculum, promoting the growth of methanogenic bacteria responsible for anaerobic digestion.

Based on the study by Wang et al. (2018), the HRT was set at 7 days, ensuring a sufficient period for the metabolic reactions necessary for the conversion of organic matter into biogas. Additionally, the medium was renewed at a rate of 250 mL per day, maintaining system balance by replacing part of the digestate with fresh organic matter without significantly altering internal conditions.

The biogas generated was measured daily using a combustible gas detector, which provided the concentration of combustible gases (in ppm). Furthermore, following the approach of Vivallos Soto et al. (2022), the total volume of biogas produced was collected and recorded (in mL), allowing precise quantification of the system’s daily production. This approach combined qualitative and quantitative characterization of the biogas, providing key data on its composition and volumetric performance.

The experiments were conducted within the mesophilic range, which optimized microbial activity and biogas production. The detailed characterization of the inoculum, substrate, and the resulting mixture after pretreatment is shown in Table 2.

**Table 2.** Characterization of the pH and TDS conditions of the inoculum, substrate, and mixture used in the assays.

| Parameters | Inoculum | Substrate | Mixture |
| --- | --- | --- | --- |
| pH | 6.22 | 6.15 | 6.82 |
| TDS (mg/L) | 425 | 790 | 810 |

### 3.3. Analytics and Monitoring

Continuous monitoring of temperature, pH, and TDS inside the reactor was performed using monitoring sensors connected to an Arduino Mega 2560 and data collected by a Raspberry Pi 3b+. The sensors used included the Gravity: Analog pH Sensor - Meter, the KS0429 Keyestudio TDS Meter V1.0, and the AZ-Delivery DS18B20 Temperature Sensor. With this system, we were able to collect data from the biodigester continuously every half hour, enabling a more thorough analysis of its conditions.

The analytics conducted focused on using accessible instrumentation for agricultural and livestock areas in an economical and efficient manner. The emphasis was on tools and technologies that are easy and inexpensive to acquire, where the mentioned sensors detect an electrical signal, which is then converted into a digital format through processing, analyzed, and stored on a computer (Teixeira et al., 2017).

For the measurement of the pH of the inoculum and feedstock before their introduction into the bioreactor, as well as for the digestate, a pH meter 3012B was used. Biogas generation was measured using the MESTEK CGD02A detector, which records the concentration of combustible gas in ppm [mg/L].

The total dissolved solids were measured using the EC Meter LS310 Quality Tester, which records values in parts per million (ppm). This high-precision sensor is specifically designed for the accurate assessment of various environmental parameters.

The TDS measurements were conducted within a range of 0 to 2000 ppm, with a resolution of 1 ppm, ensuring precise readings. The device offers a high level of accuracy, with an error margin of ±3% full-scale for TDS, guaranteeing reliable data acquisition.

For the measurement of nitrogen, potassium, and phosphorus (NPK), colorimetric tests were carried out using a commonly used colorimetric measurement kit in agriculture. Depending on the color range produced in the test, the concentration of NPK parameters was provided in ppm.

## 4. Results and Discussion

### 4.1. Biodigester Construction

The biodigester described below was designed to be easily accessible and implementable. This design enables the operation with high organic loadings in a straightforward, simple, and cost-effective manner, making it accessible for agricultural, livestock, and rural areas.

As shown in Figure 2, the mechanism for opening the digestate outlet valve is noted for its simple yet efficient design, utilizing a low-cost faucet commonly used in irrigation systems. This solution not only reduces costs but also ensures a practical and functional operation.

**Figure 2.**
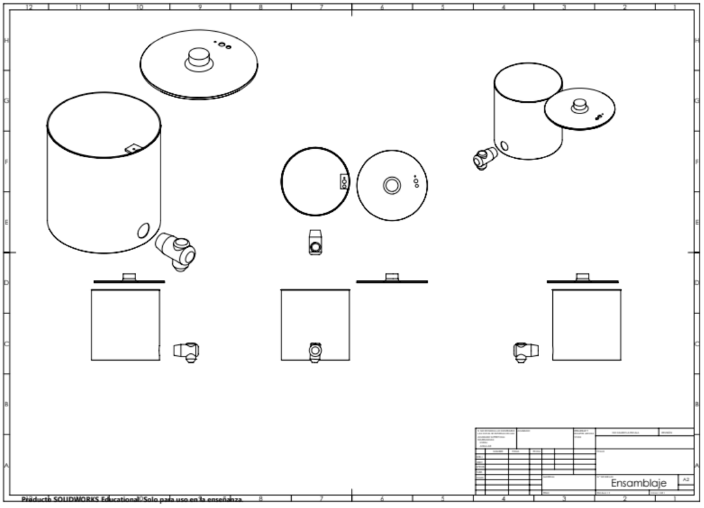
Schematic design of the assembly of each component of the biodigester, including the inlet and outlet for the feedstock and digestate, respectively.

Additionally, the cork stopper used as the opening mechanism for feeding the biodigester provides an accessible and user-friendly design, facilitating the quick and efficient incorporation of organic matter into the system.

The Figure 2 shows the holes present in the lid of the fermenter. These holes are primarily designed to allow the installation and adjustment of a hose for the outlet and venting of the biogas produced. This improvement facilitates the collection of biogas into a Tedlar bag, enabling accurate measurement of the gas volume produced during the process.

The design of this bioreactor is based on a prefabricated continuous biodigester model (Uche et al., 2020), utilizing plastic materials such as structured polyethylene (PE) and PVC. These materials not only simplify the manufacturing and assembly of the system (Vogel et al., 2023), but also offer significant economic advantages, being up to 70% cheaper than biodigesters made from metallic materials such as steel (Teixeira et al., 2017). Furthermore, the operational simplicity of biodigesters made with these plastics eliminates the need for highly specialized personnel, which favors their large-scale implementation, especially in low-cost environments (Ahmed et al., 2016).

Regarding biological performance, various studies have shown that low-cost reactors made from PE can be as effective as traditional ones in terms of biogas production, as they promote the growth of methanogenic bacteria (Garfí et al., 2016). These bacteria, essential for the anaerobic digestion process, thrive in anaerobic and thermal conditions, which promote biogas generation. It has been shown that increasing temperature accelerates the metabolism of these bacteria, especially under thermophilic conditions, where temperatures can exceed 45°C (Wang et al., 2018), significantly increasing methane production.

However, this bioreactor has been evaluated within a mesophilic range, maintaining a constant temperature of approximately 35°C, using a fish tank heater to stimulate the anaerobic digestion process. This innovative approach allows for the optimization of operating costs, maintaining adequate efficiency without the need for more expensive heating systems.

It is important to highlight that materials like PE have not been tested at high thermophilic temperatures (above 50°C) (Ahmed et al., 2016), as prolonged exposure to these conditions could lead to structural deformation of the bioreactor, affecting its performance and lifespan (Vivallos Soto et al., 2022). This limits the use of this material for applications in higher thermal ranges, which should be considered when designing systems intended to operate in environments with temperatures above 45°C (Vinnerås et al., 2006).

Another key innovation of this biodigester is the automation of the agitation system, which is carried out by a metal rod connected to a helical gear motor with a gearbox, allowing precise control of the agitation speed. This system ensures that the digestate is in constant motion, facilitating the homogeneous distribution of the components within the biodigester. Continuous agitation helps maintain constant contact between the feedstock and the inoculum, thereby promoting the microbiological and chemical reactions that occur during anaerobic digestion (Romero-Güiza et al., 2021). This agitation process also contributes to the aeration of the contents, promoting a more efficient environment for the activity of methanogenic bacteria, which in turn improves the conversion of organic matter into biogas. The ability to adjust the agitation speed allows for more precise control of the conditions within the biodigester, optimizing performance and biogas production, making this automated system an innovative and effective solution to enhance the efficiency of anaerobic digestion (Wang et al., 2018).

The remaining holes correspond to the outlets for the wiring of monitoring and sensors placed inside, which are discussed in the Section 4.2.

### 4.2. Monitoring of the Biodigester

Figures 3, 4, and 5 illustrate the evolution of temperature, pH, and TDS throughout the experimental period. Data were recorded every 30 minutes, ensuring high granularity in the results. However, periods of more linear trends are evident in Figures 4 and 5, which correspond to sensor malfunctions that resulted in data loss. For these intervals, linear interpolation was applied between the known values before and after the signal loss to impute the missing data.

**Figure 3.**
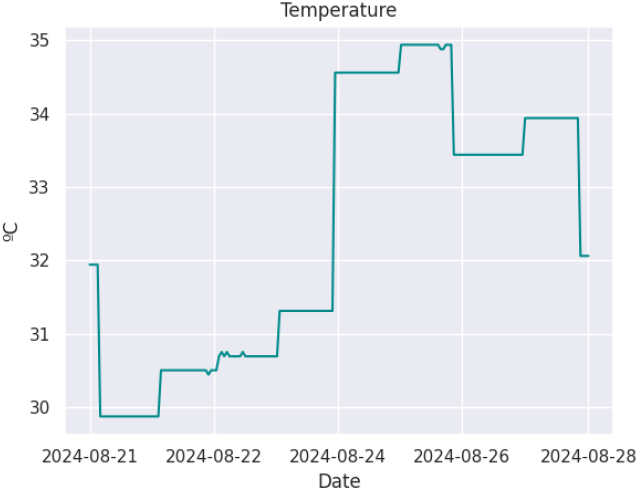
Constant temperature evolution inside the bioreactor over the 7-day testing period.

**Figure 4.**
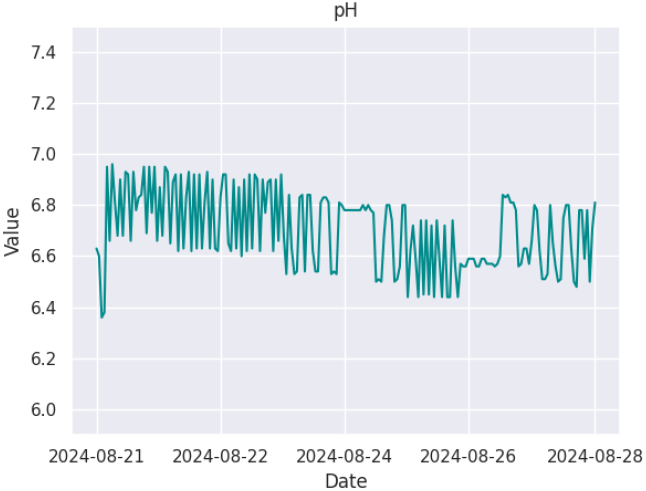
Constant pH evolution inside the bioreactor over the 7-day testing period.

**Figure 5.**
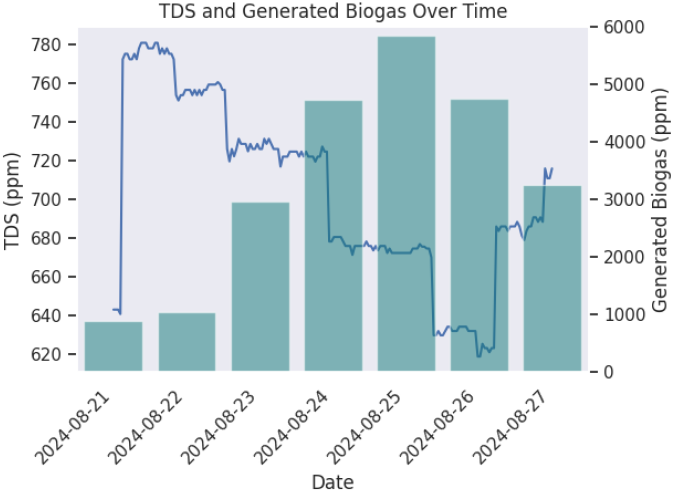
Evolution of TDS and biogas parameters over the 7-day testing period.

**Figure 6.**
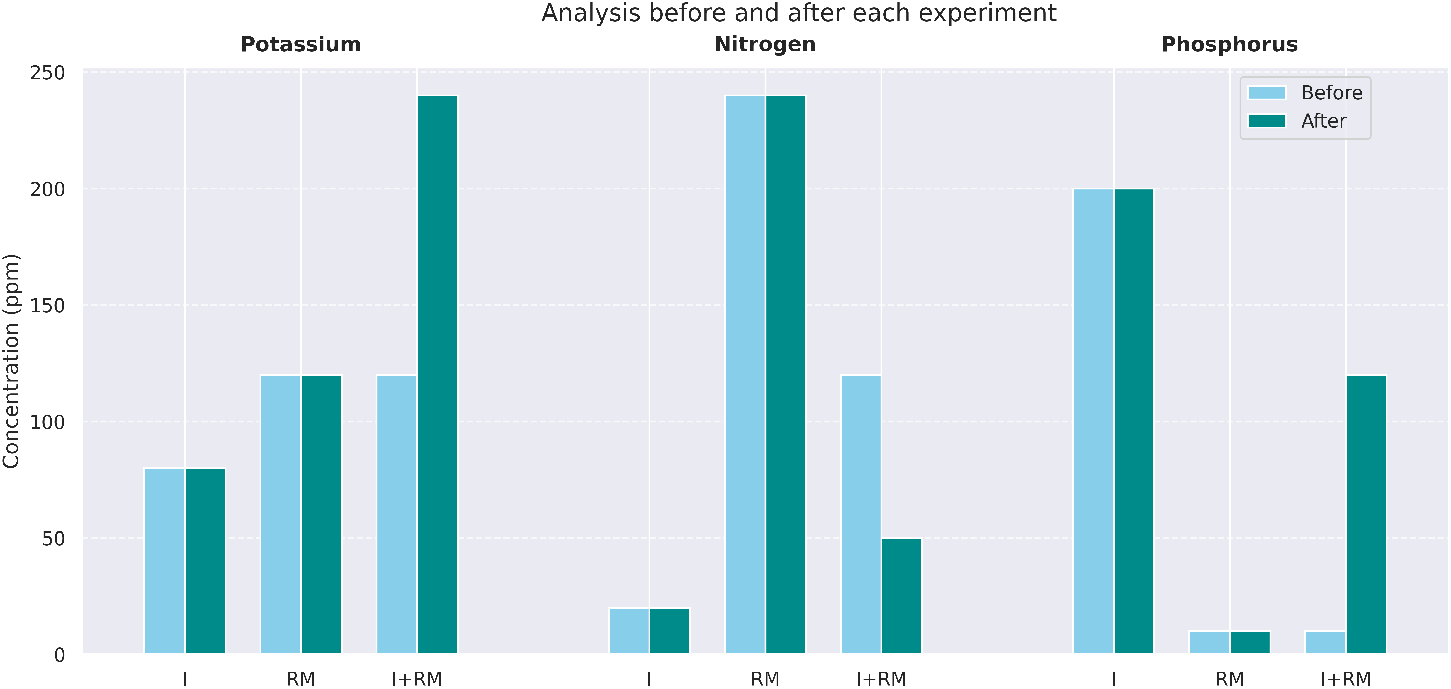
Comparison of initial and final concentrations of N, P, and K in the inoculum, substrate, and mixture after the 7-day testing period.

Following the analysis of the substrate and inoculum, as detailed in Table 2 and Section 3, and after the pretreatment process, the biodigester was loaded. The experimental conditions were previously described in Section 3.

Data were collected daily, enabling the calculation of biogas production yields during the anaerobic digestion process. This process is influenced by several operational parameters, including TDS, temperature, and pH. The evolution of these parameters inside the biodigester is presented below.

The temperature profile, as shown in the corresponding graph, ranged from 30 to 35°C, which is within the mesophilic range characteristic of methanogenic bacteria. This aligns with findings from Masse & Masse (2001), which showed the highest methanogenic activity at 30°C, followed by 25°C and 20°C. The aquarium heaters in the biodigester provided a stable environment conducive to biogas generation by maintaining an optimal temperature range for methanogens.

The pH evolution during the seven days of the experiment remained stable between 6.5 and 7. According to the study by Sivakumar et al. (2012), which investigated biogas production from spoiled milk, a pH of 7 yielded the highest biogas production.

The TDS evolution showed a maximum of 780 mg/L and a minimum of 620 mg/L, coinciding with biogas production trends. Biogas production increased to 5859 mg/L with a higher TDS, consistent with the observations of Deepanraj et al. (2015), which attribute this to the abundance of easily biodegradable materials in the substrate at the start of the trial. Biogas production evolved from an initial 887 mg/L to 3249 mg/L on the last day of the trial. However, after reaching the maximum TDS, a decrease was observed, the value on the last day of the test represents a 55% decrease in biogas production from the maximum value.

The reduced biogas production correlates with lower TDS, as inadequate microbial or enzymatic contact with the substrate occurred due to the substrate’s excess presence in the reactor. This metabolic slowdown led to a subsequent TDS increase, observed as a rise to 720 mg/L on August 28, representing a 116 % increase in TDS compared to the minimum value recorded at peak biogas production. According to Balsam & Ryan (2006), the effect of TDS on biogas production from animal waste and manure is critical, as it influences bacterial movement and growth, facilitating nutrient dissolution and transport. Water in the digester plays a key role, with total solids content directly correlating with water levels. As Sadaka & Engler (2003) highlighted, total solid concentrations below 7 % destabilize material degradation into biogas, while levels above 10 % risk overloading the fermenter.

The reduction in TDS observed between days 22 and 24 is associated with the stable pH range of 6.6 to 7. Studies by Deepanraj et al. (2015) show that reactors operating at a pH around 7 achieve higher TDS removal efficiency compared to lower pH levels. On day 26, when the minimum pH was recorded, degradation efficiency declined. Thus, maintaining a pH close to 7 resulted in better degradation efficiency compared to other pH levels.

Finally, regarding temperature, the highest increases were recorded between days 24 and 26, coinciding with the minimum TDS levels. This decrease in solids allowed for more uniform heat distribution throughout the medium. These days also corresponded to the maximum biogas production, consistent with Zhang et al. (2007), who demonstrated that upward temperature fluctuations over short periods have a more pronounced negative impact on maximum specific methanogenic activity compared to downward fluctuations.

### 4.3. Utility and Functionality of the Biodigester

In order to evaluate the functionality of the biodigester from an environmentally appropriate perspective, focusing on the reduction of harmful parameters for the environment, a study was conducted on the variability of NPK before and after anaerobic digestion.

The most common pollutants affecting water quality in the United States and Europe are NPK (Uludag-Demirer et al., 2005). Considering that this study focuses on using raw materials sourced from agricultural and livestock areas where biogas can be produced on-site, and recognizing that these areas are continuous sources of these three pollutants, these parameters were evaluated before and after AD. AD was assessed as a treatment technology for mitigating contamination at the source.

A reduction in total nitrogen (N_tot_) was observed after anaerobic digestion. The N_tot_ concentration in the inoculum was 20 mg/L, while in the raw material (horse manure), it was 230 mg/L. This nitrogen served as the basis for anaerobic biotransformation of proteins in the manure and inoculum, which were subsequently mixed in the biodigester. Proteins were first broken down into amino acids and later into ammonia, reaching a concentration of 130 mg/L. Ammonia was then used as a growth nutrient by anaerobic bacteria (Angelidaki & Ahring, 1993). This process of nitrogen compound mineralization in the organic residues, primarily proteins, resulted in forms such as ammonia (NH_3_) and ammonium (NH_4_^+^). By the end of the experiment, the final concentration of N_tot_ had decreased to 45 mg/L.

Potassium levels showed no significant change after anaerobic digestion. This is because metabolic and chemical processes during AD do not significantly affect potassium concentrations. Consequently, the potassium concentration in the reactor effluent corresponds closely to the combined concentration from the inoculum and raw material (Yoshino et al., 2003). In the specific case of horse manure, potassium tends to dissolve during anaerobic digestion and generally remains in a soluble form, with minimal chemical interactions altering its concentration. Thus, the potassium concentration in the effluent is similar to or even slightly higher than in the starting material.

Regarding phosphate, Phosphate-solubilizing microorganisms can release soluble inorganic phosphorus during the decomposition of organic compounds. This phosphorus can either be utilized by microorganisms or released into the effluent (Münch & Barr, 2001). In this study, the inoculum initially contained 200 mg/L of phosphorus, but the effluent exhibited a 65 % reduction in concentration. Most anaerobic microorganisms lack the capacity to mineralize phosphorus at large scales, as noted by Yoshino et al. (2003), the reduction in soluble phosphorus can be attributed to elevated pH conditions (approximately 7 in this study) or the presence of high concentrations of cations such as calcium, magnesium, or iron. These conditions promote the precipitation of phosphorus into insoluble forms, such as apatite (Ca_5_(PO_4_)_3_OH) or struvite (MgNH_4_PO_4_·6H_2_O). This process is particularly significant in systems with excess ammonia, as observed in the nitrogen data.

As observed, after the anaerobic digestion process, the designed bioreactor has proven to be an effective environmentally sustainable solution, significantly reducing total nitrogen and phosphorus present in the input feedstock. Consequently, the resulting digestate exhibits lower NPK concentrations, contributing to a reduced environmental impact.

## 5. Conclusions

The design and construction of the homemade bioreactor developed in this study proved to be an effective tool for biogas production from organic waste, particularly horse manure, in rural and agricultural settings. The monitored parameters—pH, temperature, and TDS—played a crucial role in the efficiency of anaerobic digestion by promoting methanogenic activity. A pH near 7 and mesophilic temperature range of 30–35°C supported optimal biogas yields, while TDS fluctuations significantly influenced production. High solid concentrations did not always enhance efficiency and, in some cases, reduced it due to fermenter overload.

Regarding contaminants, a significant reduction in total nitrogen was observed, from 230 mg/L to 45 mg/L, driven by the biotransformation of proteins into ammonia and its subsequent utilization by anaerobic bacteria. Potassium, in contrast, remained largely unchanged, as it predominantly exists in a soluble form during anaerobic digestion. Phosphorus concentrations decreased by 65 %, likely due to precipitation into insoluble forms facilitated by elevated pH and the presence of cations such as calcium, magnesium, and iron.

This study highlights the viability of anaerobic digestion as a sustainable and efficient technology for treating organic waste in regions with abundant biodegradable materials. Additionally, the results demonstrate the potential of this process not only for biogas generation but also for reducing contaminants in effluents, contributing to a more efficient and environmentally friendly waste management cycle.

## CRediT authorship contribution statement

**Ángela Ventura-Prados:** Conceptualization, Methodology, Validation, Investigation, Resources, Writing - Original Draft, Project administration. **Javier Solís-García:** Software, Data Curation, Formal analysis, Visualization, Writing - Review & Editing.

## Notes

### Competing Interest Statement

The authors have declared no competing interest.

